# Tracing the genetic etiology of cardiovascular disease using a hierarchy of common genetic variants derived from patient subgroups stratified by differential levels in severity

**DOI:** 10.1101/213124

**Authors:** Sarosh N. Fatakia

## Abstract

A pedagogical review in informatics based on computational molecular evolution is synopsized to illustrate the role of common genetic variants during adaptation to diverse habitat and physiological functions across disparate species. With that background, an evolutionary perspective is motivated to trace lifestyle-related disease progression in humans. Cardiovascular disease (CVD) is a multi-factorial disease, where maladaptation due to a sedentary lifestyle and faulty diet may influence its prognosis, but a healthy diet and lifestyle may as well restrict it. However, a comprehensive genetic basis for the differential severity in CVD remains unreported. Here, we have computed that CVD severity may be associated with the hierarchical plasticity of genes whose common genetic variants are exclusive among those patients having disparate levels of disease severity. Most importantly, we have used a small and outbred subpopulation to demonstrate that constellation of common variants can be exploited to trace the disease etiology. Taken together, we report a novel perspective to identify a hierarchy of common variants associated with differential CVD severity and subsequently hypothesize how the constellations of common variants may collectively modulate disease severity, which in turn may result in a relatively broad and complex spectrum of severity at the population-level. To corroborate our findings, we report a hierarchical plasticity of *LDLR* gene, which has been previously implicated in the differential response to lipid metabolism, is associated with differential CVD severity in the subpopulation under consideration.

## Introduction

Human evolution [1-4] is a continuous process of adaptation [5, 6] during which we have developed immunity against diseases in general [7], as well as local to a region [8], and also kept pace with the dietary changes [9], external conditions of habitat [10-12], and geography [13]. In a comparative genomic study, we may invoke conserved (or invariant) genetic information to trace the shared ancestry [14, 15]. Inspired by a recent report that a healthy lifestyle may reverse some adverse consequences related to cardiovascular disease (CVD) [16], and merging comparative genomics with evolutionary biology, we present insights and new perspectives to rationalize the disease etiology. Moreover, it has been long been suggested that an evolutionary standpoint may improve our understanding of common lifestyle-related diseases such as CVD [17, 18], and recent reports suggest the limitations of converging toward a consensus set of causative variants from diverse studies [19-22].

Cardiovascular disease has been attributed to a plethora of genetic variants, which involve lipid regulation [23-26], inflammation [27-29], as well as other epigenetic agents such as an altered lifestyle [30, 31]. Genome wide association studies involving thousands of individuals, have identified hundreds of causative and associated variants to understand the genetic etiology of CVD [32-35], but it has been estimated that the spectrum of identified variants does not fully represent the inherited risk [22, 32-36]. The genetic etiology of such lifestyle-related diseases can seldom be identical in an outbred population because their ancestries may differ [37]. Hence, not all variants may be identified with high statistical confidence from a specific genome wide association study (GWAS), as suggested schematically in Fig. 1 (adapted from [38]). Therefore, most importantly, results of GWA studies, from distinct subpopulations may differ because the causative variants may not be identified with similar statistical confidence and the ancestry of individuals with CVD may also differ [36].

**Figure 1.**
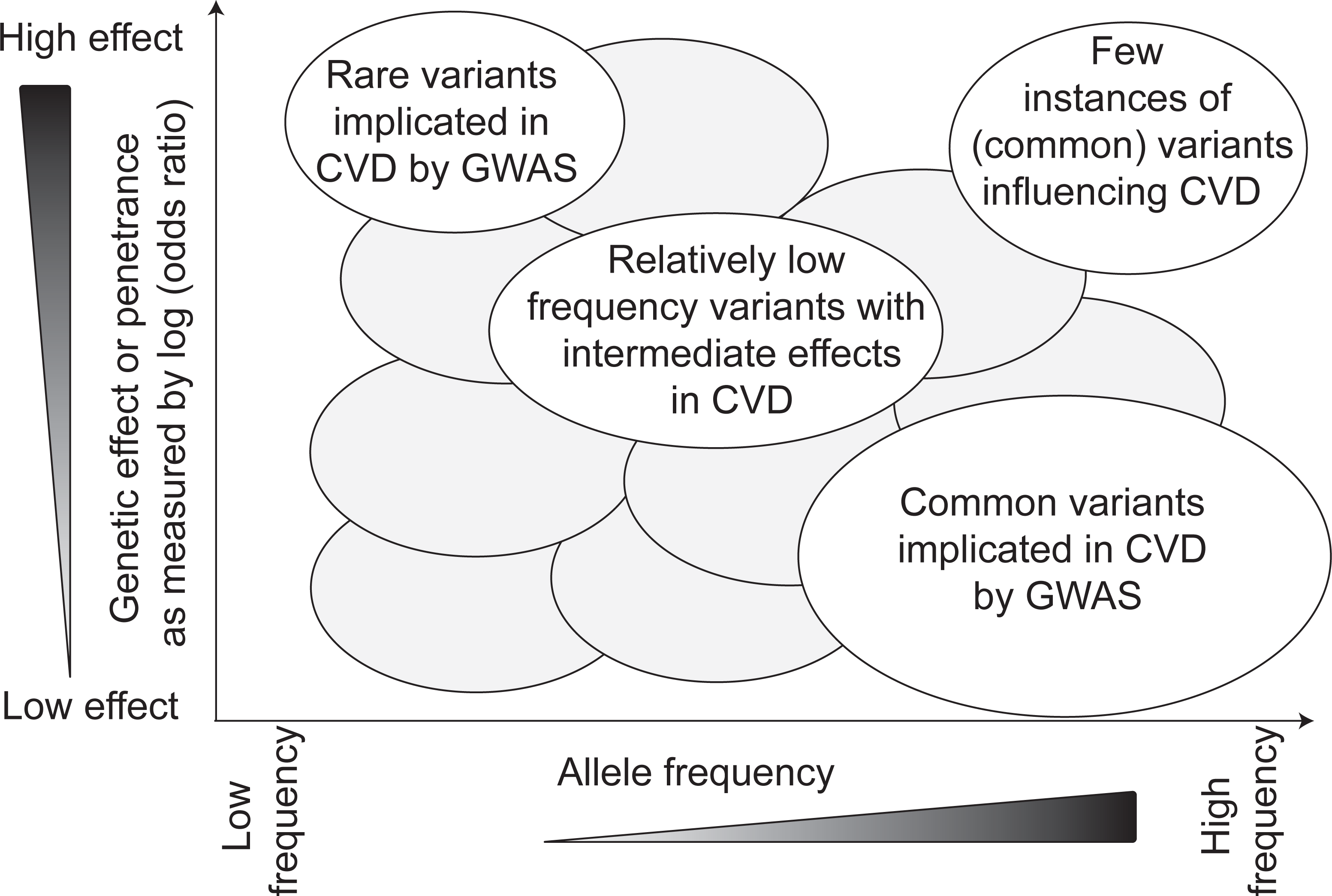
A schema reveals the trend for volume of data needed to characterize the genetic etiology of a multigenic disease. Shaded hypothetical clouds illustrate a continuum of values of allele frequency, which may range from the low end of the spectrum (lighter shade of triangle) to the high end of the spectrum (darker shade of triangle). The region of interest probed from low volume data studies may only pertain to identification of common variants, which are considered causative for relatively low penetrance (bottom right corner, or middle right region of graph). GWAS = genome wide association study.

Besides a well-documented set of candidate genes associated with cardiac diseases [39], there are other genes that may differentially affect their prognosis, for example DNA damage repair (DDR) genes *TP53* and *MDM2* have been recently implicated in cardiomyocyte function [40]. In this report, we propose that an evolutionary study may be invoked to trace a genetic etiology of CVD in an outbred subpopulation, and overcome the challenges of a small data set. From our proposed formulation, we may be able to corroborate previously implicated CVD genes (and their variants) and also identify new ones. As onset of CVD is largely lifestyle-related in the general population [16], we hypothesize that the disease etiology may be traced by contrasting distinct variants of the same gene. In fact it is well established that specific mutations (generally referred to as variants) of the receptor LDL receptor gene (*LDLR*) cause hypercholesterolemia, and might lead to high risk of myocardial infarction [41], but other mutations of *LDLR* may in fact be protective against increased LDL levels [41].

Using microarray technology, an invariant genotype can be detected from the genome-wide genotype that was probed across various individuals in an outbred subpopulation, at a specifically predetermined single nucleotide polymorphic (SNP) locus. In a comparative genomic study across those subjects, the genotype for a hypothetical common ancestor for the specific cohort may thus be unambiguously determined at that SNP locus. Their hypothetical last common ancestral genome of that outbred subpopulation will also manifest the same invariant genotype at the SNP locus, which has remained invariant subsequent to evolution [42]. Hence, we hypothesize that such SNP loci with invariant genotype (common alleles from that subpopulation), which have resisted mutations, may be utilized to trace the genetic etiology of CVD cases with respect to age- and gender-matched controls. The occurrence of common genetic variants at any given genomic locus is an inevitable consequence of evolution within an outbred population [43], because those variants may progressively get inherited over generations [44]. Such common variants are likely to be a relatively recent outcome of evolution compared to invariant ones, but presumably older than the relatively rare variants [44]. However, there is a caveat that rare variations are either likely to be new (few generations old) and hence they have not been subject to negative selection for a long time, or are rare because they are being selected against owing to their deleterious genetic nature and are likely have greater penetrance in contrast to that due to common variants [44-46].

By partitioning human noncoding sequence into invariant (conserved) and variant (non-conserved) loci, it has been reported that agents of natural selection have influenced the human genome evolution [47]. Hence, from the perspective of a comparative genomic approach, interactions involving both invariant and variant genomic loci may lead to gain-of-function (adaptation) or loss-of-function [48, 49] or perhaps remain benign without any perceptible change at the genome-level [50, 51]. However, the adverse consequences of these changes may manifest later on during mid-life, much beyond the average reproductive age at the population level. For example, it has been suggested that high levels of cytokine response may confer an innate resistance to infectious diseases at an early age, and that the same response inadvertently becomes adversely associated with CVD at a much later age [52]. An evolutionary adaptation that confers reproducibility success at a relatively early age, but which may adversely affects us later, has also been reported with regard to an ancient variant allele at the growth differentiation factor-5 (*GDF5*) enhancer region [13]. Taken together, the constellation of genetic variants may have a profound impact at the individual genome level [53-55]. Therefore, by reaching an appreciable frequency in a subpopulation, common variants (evolutionary adaptations) are instrumental for a differential physiological response such that the level of severity may be stratified. Therefore, we propose that the genetic etiology for CVD may be investigated in an outbred subpopulation of patients and age-matched healthy controls using common variants identified from patients with stratified severity level of this disease. We hypothesize that a hierarchical nature of those common genetic variants among patients with disparate severity may help trace the evolution of CVD severity.

Traditional GWA studies for complex multifactorial diseases warrant genome-wide data from thousands of individuals to identify causative variants with high statistical confidence [38]. Therefore, it has been a challenge to characterize the genetic etiology of CVD from India with access to genotype information from nearly one hundred and fifty CVD cases, and an identical number of age- and gender-matched controls. To overcome the limitation of a small data size, we propose insights and perspectives for an inter-disciplinary approach, and hierarchically trace genome-wide common genetic variants among CVD cases with stratified levels of severity that was determined from angiography-based studies [56, 57].

At first, we must present a primer on protein sequence evolution to elucidate its potential using published reports. This synopsis relates to proteins in general, and DNA damage repair proteins in particular, but the insights developed here are for a relatively wider and general application to genomes. Therefore, the insights and perspectives here are addressed to the larger community, inclusive of clinicians and non-clinicians.

## Genome sequence evolution using computer-based comparative genomics approaches

### The DNA damage repair genes

All eukaryotic living cells experience DNA breaks in their nucleus due to a plethora of endogenous and exogenous agents, and have evolved intrinsic molecular mechanisms to repair those lesions [58]. If left unrepaired, the physiological consequences may be lethal for the cell [59]. Such mechanisms for repair constitute a network of bio-chemical pathways called DNA damage response (DDR) pathways [60], the regulation of which is largely evolutionarily conserved [61]. For example, in the instance of metabolic syndrome, it has been suggested that system-level DDR may impinge on energy metabolism and vascular physiology [60]. Moreover, when the genes that encode human DDR proteins undergo mutations, the inherent genome-wide repair mechanism may breakdown, which may lead to ischemic stress [62], CVD [63, 64], or certain cancers [65-67]. Over a hundred different genes encoding the various human DDR proteins [68, 69], and belonging to disparate DDR pathways [58, 70], are publicly available at: http://sciencepark.mdanderson.org/labs/wood/dna_repair_genes.html.

Evolutionarily related proteins (or the genes encoding them) are referred to as homologs. The homologs within the same organism, which are a consequence of gene sequences that coalesce [71] at specific within-genome replication event (such as a gene duplication event) are termed paralogs. On the other hand, when gene sequences coalesce [71] at a speciation event they give rise to orthologs. The mathematical formulations of this coalescent theory have been extensively implemented (for example [72]) and covered extensively (for example [73]). Genes encoding orthologs and paralogs have largely evolved by gene duplication events, as well as other evolutionary processes such as retroposition, intron-mediated recombination (exon shuffling and trans-splicing), horizontal gene transfer and de novo organization (reviewed in [74]). For this comparative genomics approach, an accurate multiple sequence alignment (MSA) is critical for drawing an accurate inference [75]. The MSA is primarily a mathematically derived arrangement of nucleotide or amino acid (AA) sequence (or may merely represent domains / motifs / fragments of the full sequence), of three or more homologs representing their overall ancestry.

Homologous proteins have evolved among diverse species, and the ones that are critical for multiple functions, may largely remain conserved at the amino acid residue (AA) level, and under strong negative selection [76-79]. As most DDR proteins are ubiquitous and play a relatively central role in cellular processes within an organism, it is plausible that they are largely under strong selection pressure and evolutionarily conserved [78, 80]. However, random mutations can give rise to variants at the codon-level, which in may turn out advantageous and may get incorporated or “fixed” in a population, or subpopulation, by natural selection [81]. From among all possible mutations, the ones that are advantageous to the organism’s fitness, and enable it to adapt and thrive in its habitat will be discussed later. These genomic adaptations are not exclusively single nucleotide variants (SNVs), and include instances when molecular adaptions in host organisms were geared to avert insertion of mobile elements, such as transposons, from parasite genomes [82, 83]. However, before a rare mutation gets “fixed” in the population, they first manifest as polymorphisms in subpopulations, and can have interesting physiological consequences [84]. To showcase the importance of genetic invariants and common variants during genome evolution, we synopsize some select reports of molecular adaptations of among various DDR genes, across different species. To predict the AAs that are important for the protein structure and/or function, computational methods are used to delineate evolutionarily conserved AA positions in a MSA (columns of AAs). DNA repair proteins from *E. coli* and S. *cerevisiae* have been compared to protein sequences encoded in completely sequenced genomes of disparate bacteria, archaea and eukaryotes [85]. In that study, the authors identified conserved domains in DDR proteins across kingdoms and highlighted that horizontal gene transfer and epigenetic regulation influenced their sequence composition [85]. Subsequently, independent studies have identified homologous DDR proteins in disparate species, such as *E. coli, S. cervevisiae, M. musculus* and *H. sapiens* [61, 86, 87]. Moreover, it is noteworthy that analogous roles of orthologous proteins (such as RecA/RAD51) have been traced across eukaryotes: in bacteria, yeast and mammals [61, 87]. As another example, nearly two decades ago, two independent comparative sequence analysis of yeast DDR protein with its orthologs, and *E. coli* DDR protein with its orthologs have led to discoveries of novel orthologous human DDR proteins [88, 89].

An important component of comparative sequence analysis is the *in silico* modeling of the AA sequence evolution. Codon-based models of gene evolution, such as Phylogenetic Analysis by Maximum Likelihood (PAML) [90] and Selecton [91] (see Box 1), use MSAs of closely related nucleotide sequences to compute the evolutionary selection pressure at the codon-level, and differentiate positive selection with respect to negative selection and neutral evolution.

#### Box 1

Special codon-based models of gene evolution, such as modules/suites available from PAML [90] and Selecton [91], have been developed to simulate single nucleotide substitutions among orthologs and paralogs. Among the various modules implemented, the one most relevant to our report is the one that simulates SNVs to compute the evolutionary selection pressure at the codon-level for a given MSA of homologs. “Silent” mutations, which are synonymous nucleotide substitutions do not change the translated AA sequence so their substitution rate *d_s_* (also called Ks) is not subject to a selection pressure on the expressed protein [80, 92]. Such mutations are more prevalent in an organism compared to the nonsynonymous ones. The nonsynonymous substitutions alter the amino acid sequence and their substitution rate *d_N_* (also called *K_A_*) is a function of the selective pressure on the protein [80, 92]. A measure of the selection pressure at the codon level is subsequently obtained from their ratio 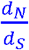 (also called 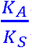), also referred to as [80, 92]. To ensure an unbiased estimate of this ratio, an average of one nonsynonymous substitution may be faithfully represented at the codon level (numerator *d_N_* ~ l). If relatively distant sequences are compared, the computation may be biased, as there is ambiguity with regard to whether only a unique single nucleotide substitution, or multiple such substitutions were accrued at the codon level. Hence, if we use distant orthologs, there is a likelihood that *d_N_* > 1 and may not be faithfully represented [93]. Caveat: an estimate of *d_N_* may be computed at first, before computing, using the same suite of software packages [90, 91].

### Differential changes in the AA protein sequence across diverse species

Computational sequence analysis has made it feasible to identify homologous protein sequence, and subsequently narrow down the evolutionary adaptations (or variations) in those genes. Therefore, one may use a candidate gene-based approach to study select proteins (and their homologs) along with their interacting proteins (dubbed interactors). Here, we have highlighted one such DDR protein, p53, which has been extensively studied, to provide a wider perspective (See Box 2), but in theory any set of proteins may be analogously showcased.

#### Box 2

##### P53 as a representative example for comparative study across species

The human tumor suppressor gene (*TP53*) (an oncogene) encodes for the tumor suppressor protein 53, referred to as P53. To showcase its multiple roles in fundamental cellular functions, it is referred to as the “Guardian of the Genome” by David Lane [94], and as the “Cellular Gatekeeper” by Arnold J. Levine [95]. This protein and its homologs have been extensively studied in regard to critical cellular processes such as cell cycle and DDR pathway [96]. Therefore, we synopsized molecular adaptations of P53 homologs and a select subset of its interacting proteins (subsequently referred to as a p53 sub-interactome).

##### A primer on p53 structural domains

A schematic of the various P53 functional domains is represented as a 1D cartoon in Fig. 2 (Adapted from [97]). The N-terminal contains a short intrinsically disordered region, following that is the transactivation domain (TAD), which is further divided into two subdomains (TAD1 and TAD2) that bind to negative regulators, transcriptional coactivators and components of transcriptional machinery [98-101]. Subsequent to that is a Proline-rich region (PRR) that favors a polyproline II helix structure and serves as a rigid linker to project the TAD distal to the DNA binding domain (DBD) [102]. In contrast to the TAD, DBD is more conserved across disparate vertebrates. Following the DBD is the C-terminal domain (CTD), comprising the tetramerization domain (TET) and the extreme C-terminus (CT).

**Figure 2.**
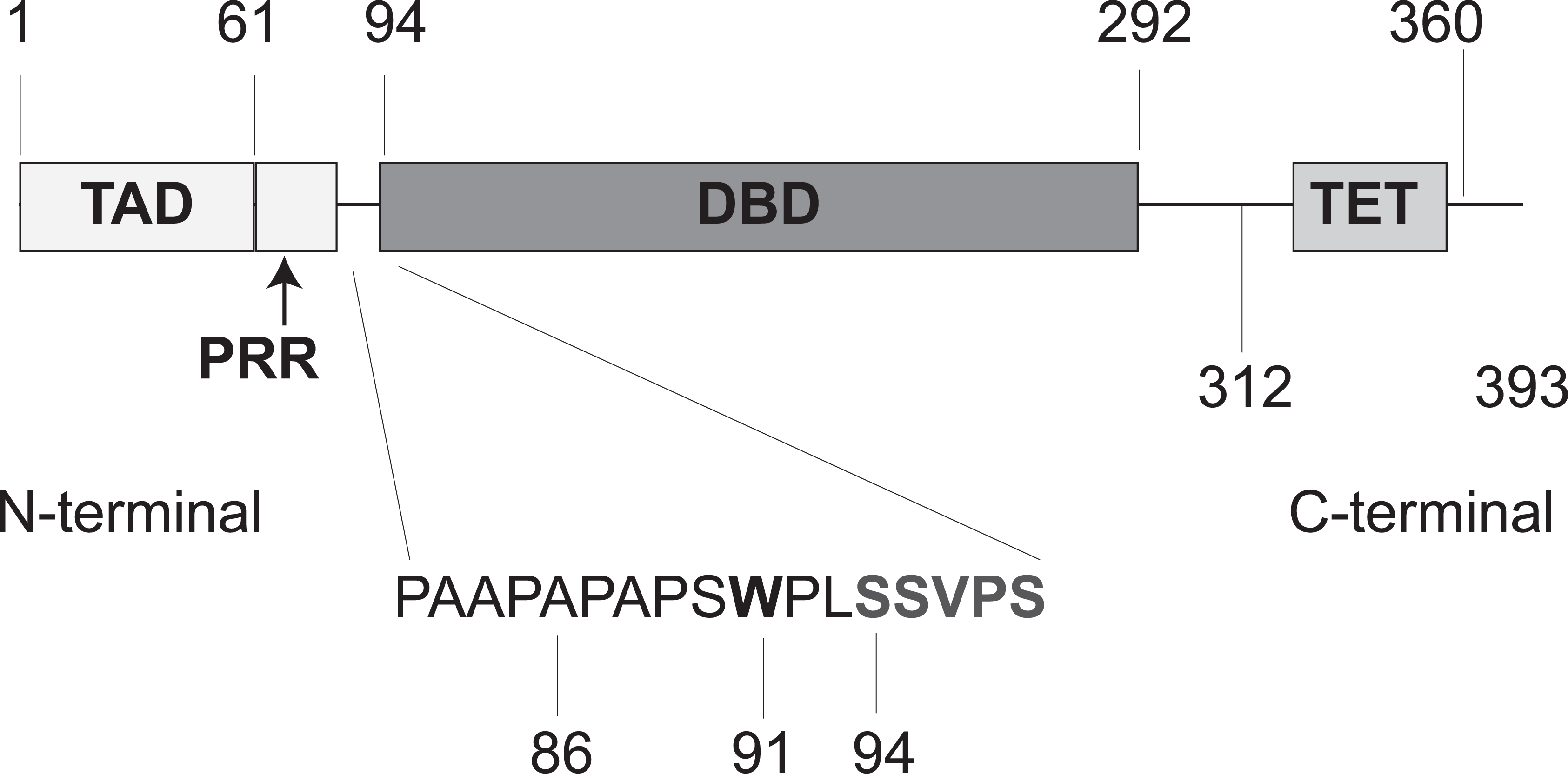
The functional domains from the p53 protein sequence are described. The human p53 molecule comprises of 393 amino acid residues, whose positions ID are sequentially illustrated on the top and below the schematic. The N-terminal comprises of a transactivation domain (TAD) and proline-rich region (PRR). It is followed by the DNA binding domain (DBD), and subsequently followed by the C-terminal. (Adapted from Reference [97]). **(Inset for Box 1.)**

##### The p53 sub-interactome

Coordinated AA interactions are imperative for protein function such as folding, unfolding, and binding to other identical or non-identical proteins to form complexes. For example, it has been demonstrated that the stability of the p53 tetramer is enhanced due to specific AA interactions of the p53 DBD with the p53 N-terminal [97]. The AA critical for such essential function may manifest as an invariant AA across various orthologs from disparate species and resist mutations. Such a conserved position (or column) usually stands out when viewed in a fragmented MSA (or full-length MSA of the protein).

Using a candidate gene approach, we considered two molecular interactors of p53: MDM2 and MDMX (aliased MDM4), the genes of which have also been reported as oncogenes, being expressed in some types of cancers [103]. Murine p53 protein interacts with Mdm2 (murine ortholog of MDM2) via its N-terminal as well as core domain [104]. Moreover, it is reported that Mdm2 binds directly to the p53 TAD1 domain to inhibit p53 interactions with its transcriptional coactivators [105] and that their binding is differentially decreased when the murine p53 CTD is deleted, mutated or even acetylated [106]. In addition, the MDMX protein is an important negative regulator of p53 in normal cells. Bista *et al.* [107] have shown that the full-length MDMX contains a sequence of regulatory element (the “WWW element”) that binds to its own N-terminal domain and therefore blocks the binding of MDMX to p53, and Su *et al.* have shown that there exists a motif ‘FXXLW’ motif, which is the MDM2 functional motif, in the disordered region of p53 [108]. Such intricate molecular adaptations of these interacting proteins are critical as an inhibition introduces a level of functional regulation making them ideal candidates for study of a p53 sub-interactome with regard to their differential levels in function and protein sequence evolution. On the other hand, the p53 C-terminal binds to RAD51 and RAD54 and additional details of specific interactions are described in Ref [109, 110].

We now synopsize studies that have identified genetic variations as molecular adaptations among orthologs genes of p53 and its sub-interactome. Comparative study of orthologous genes from mammals such as primates, bats and rodents are used here to illustrate how computational biology can help identify codons under positive selection, following which experiments were used to validate those adaptations (A synopsis is presented in Box 3).

#### Box 3

##### Synopsis of species-specific adaptations in DDR proteins

We reiterate that the mechanism to repair endogenous and exogenous DNA damages have evolved across disparate species for their survival and proliferation. To maintain a robust DNA-repair pathway, it is plausible that a large fraction of genes are conserved but only a small subset of genes may be under positive selection pressure [78, 80]. Positive selection at two or more unique codons within the same gene and across genes may suggest improvement or modulation of a pre-existing function [111], for example pain-related stimuli in humans [112]. Mutations that tentatively enhance the organism’s fitness are referred to as molecular adaptations and the codons within the same gene, or across different genes, are set to have co-evolved (reviewed in [113]). In this Box, we highlight some preselected reports that have used comparative genomics to identify specific molecular adaptations among DDR proteins across species. We sought out how have the DDR genes had adapted across those diverse species that inhabited harsh and diverse environments, which were not conducive for most mammals. For example, amongst mammals, (i) bats are the only species with flight as their mode of transport, on the other hand; (ii) blind mole rats live in subterranean conditions in extremely cold and hypoxic conditions, where sunlight may not permeate.

###### 1 – Bats

The bat is the only mammal with an innate ability for sustained flight, and therefore a comparative genomic approach with respect to other diverse mammals helps uncover the putative molecular adaptations in select genes. It has been reported that nearly 23% and 5% of bat specific mitochondrial genes and nuclear encoded oxidative <u>phos</u>phorylation (OXPHOS) genes are under significant positive selection respectively [114]. This prompted further investigations to test the hypothesis that positive selection of the genes facilitated in metabolic and energy needs for a sustained flight [114]. Their physical act of flight is perhaps one of the most energy consuming physical activity and therefore it is was suggested that its genes involving energy metabolism and DDR may have evolved unique molecular adaptations, which are absent in all other mammals [115]. Next, to investigate the molecular sequence evolution of the various bat genomes, a comparative genomic approach was used to study genomes from two bat species: *Myotis davidii (M. davidii)* and *Pteropus alecto (P. alecto)*, and compare them with diverse mammals including *Homo sapiens* [115]. In that study, Zhang *et al.* delineated genetic variants from selected mammals for p53 functional domains, and other DDR genes such as *MDM2, ATM, RAD50* and *KU80* [115]. Here, in Fig. 3 (adapted from [115]), we showcase a fragment of p53 MSA, from disparate mammals including *M. davidii* and *P. alecto*. Each of the conserved AA at the MSA position suggests zero tolerance to nonsynonymous substitutions, and the rare plasticity of non-conserved positions revealed tolerance to genetic variations, suggesting putative candidates for molecular adaptations. In particular, the authors identified the K321M non-synonymous mutation as the *TP53* molecular adaptation in *M. davidii* and *P. alecto*, but the L323A mutation as an exclusive adaptation in the *M. davidii* sequence [115]. In gene-based studies, they established that the *p53* and *BRCA2* orthologs of were under positive selection pressure in *M. davidii* and that the *LIG4* ortholog was under positive selection pressure in *P. alecto* [115]. Therefore, it is plausible that the molecular adaptations of *M. davidii* and *P. alecto* occurred prior to the divergence from their last common ancestor, within the bat lineage, and the unique species specific adaptations (for example L323A in *TP53* from *M. davidii* and the adaptation in *LIG4* ortholog from *P. alecto*) occurred subsequent to that divergence [115]. In addition, Zhang *et al.* also reported that the human orthologs of *ATM, RAD50, KU80* and *MDM2* genes are under positive selection pressure, amply suggesting that molecular adaptations within a species usually brings about a concerted genome evolution [115]. We also reiterate that a mutation that impacts fitness (positive or negative) can be fixed by drift as well as selection [22]. A caveat is in order here that independent experimental assays performed in that study, suggest that those putative variants were a result of species adaptation and that the previously mentioned theoretical studies provided insights for an initial working hypothesis [115]. Moreover, experimental assays usually measure fitness consequence in terms of survival probability, and usually not in a complete physical and genomic environment “experienced” by the ancestor. Hence, in most instances, it is generally not possible to experimentally “confirm” a molecular adaptation.

**Figure 3.**
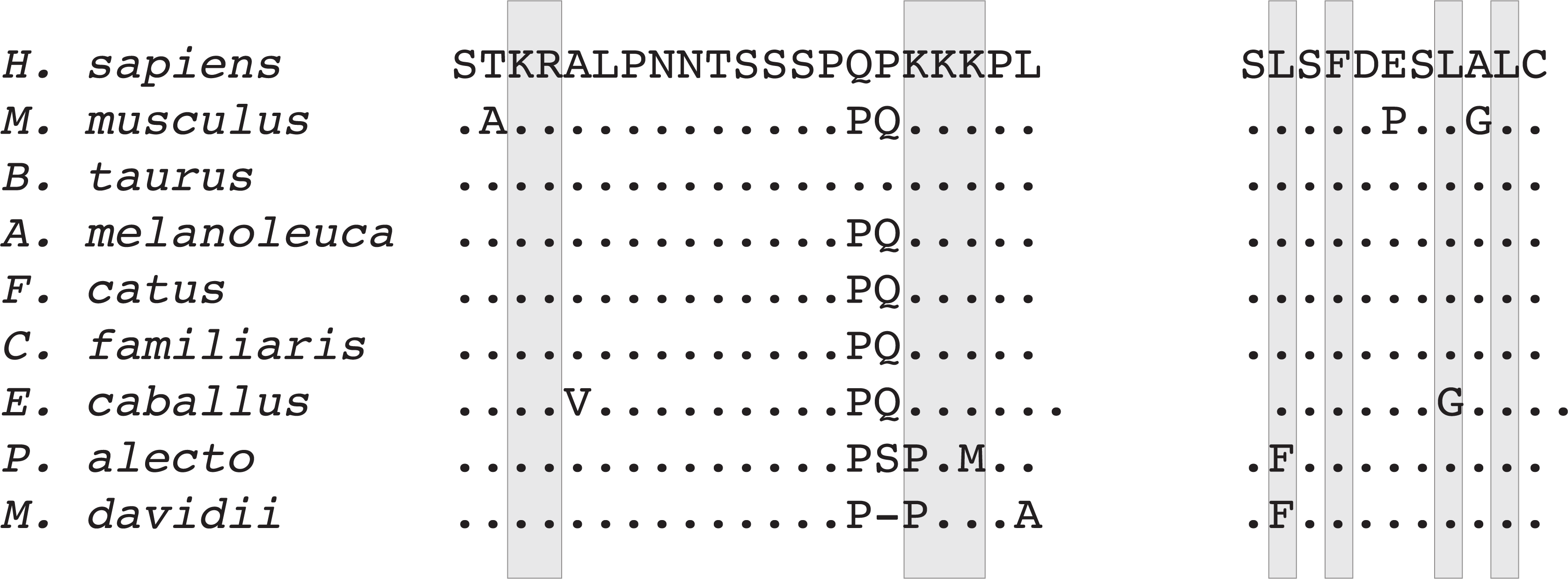
Representative MSA of p53 and MDM2 orthologs showing the different AA variants unique to bats. These AA variants were detected in the functionally relevant regions of the p53 nuclear localization signal and MDM2 nuclear export signal (shaded). The p53 MSA (shown on left) is representative MSA from nuclear localization signal. MDM2 MSA (shown on right) is representative for nuclear export signal. The dot connotes AA identical to the one in the first row of the fragmented MSA (Adapted from [115]). **(Inset for Box 2)**

Molecular adaptations among rodent species are synopsized in modules (2) and (3) below, and contrasted with other mammals.

###### 2 – Zokor rats

Zokor rats, blind subterranean rats, root rats and bamboo rats are spalacid rodents that are collectively referred to as mole-rats. These rodents thrive under adverse geographical conditions, for example the Tibetan plateau is a natural habitat for zokor rodents [116]. Recently, two subterranean wild zokor species (highland-dwelling *Myospalax baileyi* and lowland-dwelling *Myospalax cansus*), and one highland-dwelling aboveground species of root vole (*Microtus oeconomus*) were investigated to explore the molecular adaptations in the p53 and contrast them with functional adaptations related to the harsh natural habitat and environmental stress [116].

As previously mentioned, a part of the p53 N-terminal is intrinsically disordered [104], and its MSA with other orthologs represents a high degree of sequence non-conservation [116]. Zhao *et al.* first used cues from protein sequence comparison and computational biology to narrow down the search for putative molecular adaptations at the codon-level in the p53 DBD, which was subsequently cross-checked using site-specific mutagenesis assay [116]. The DBD is relatively more conserved compared to the TAD and CTD and an unambiguous MSA from DBD was first obtained across orthologs, from humans to many different rodents [116]. For the purpose of this synopsis, we highlight that the sequence conservation of domain is disrupted by a solitary non-conserved MSA position, at codon 104 in human p53 DBD. Nonsynonymous variants of p53 at codon 104 were identified in the three rodent species [116]. *P53* orthologs in primates such as *Homo sapiens, Macaca fuscata, Macaca muleta* have a Serine (S) residue at codon 104. However, *Myospalax baileyi* and other mammals such as *Ovis aries* (sheep), *Bos taurus* (cow), *Rattus norvegicus* (rat) and *Mus musculus* (mouse) have Asparagine at that corresponding homologous codon [116]. In addition, the authors reported that for *Myospalax baileyi*, the 104N variant is responsible for the transactivation of apoptotic genes at extremely severe environmental conditions: hypoxia, hypercapnia (acidic stress, high CO_2_) and severely low ambient temperatures [116]. Subsequently, they have also reported that the 104E nonsynonymous variant in p53 for *Microtus oeconomus* suppresses apoptotic gene reactivation and cell apoptosis [116].

###### 3 – *Spalax* rodent

*Spalax* a blind subterranean mole-rat belongs to the Spalacidae family. As shown in Fig. 4, the Arginine residue is conserved across p53 sequences in humans, primates, quadrupeds and numerous rodent species [117]. Comparative sequence analysis of the p53 DBD sequence among various rodent species revealed that the *Spalax*, had acquired the R174K mutation [117]. However, in addition to *Spalax*, the R174K mutation is identified across other species such as *Xenopus laevis, Monodelphis domestica, Xiphophorus maculatus*, and *Xiphophorus helleri* (Fig. 4, adapted from [117]). The above nonsynonymous substitution is one of the few mutations that was adapted within the *Spalax* genome, and the most interesting feature is that this particular nonsynonymous substitution occurs at an otherwise well conserved position in the MSA. Additional Spalax-specific changes, with respect to human and mice, were also reported using a full-length MSA [117]. All findings motivated the authors to hypothesize that Spalax *p53* gene had adapted, and enabled those rodents to thrive in an underground hypoxic environment. Subsequently, other critical adaptations related to hypoxia were also reported in *Spalax* [118]. (More recently, unique hypoxia-based adaptations have also been reported in the case of the naked-mole rat, another member of the same Spalacidae family [119]).

**Figure 4.**
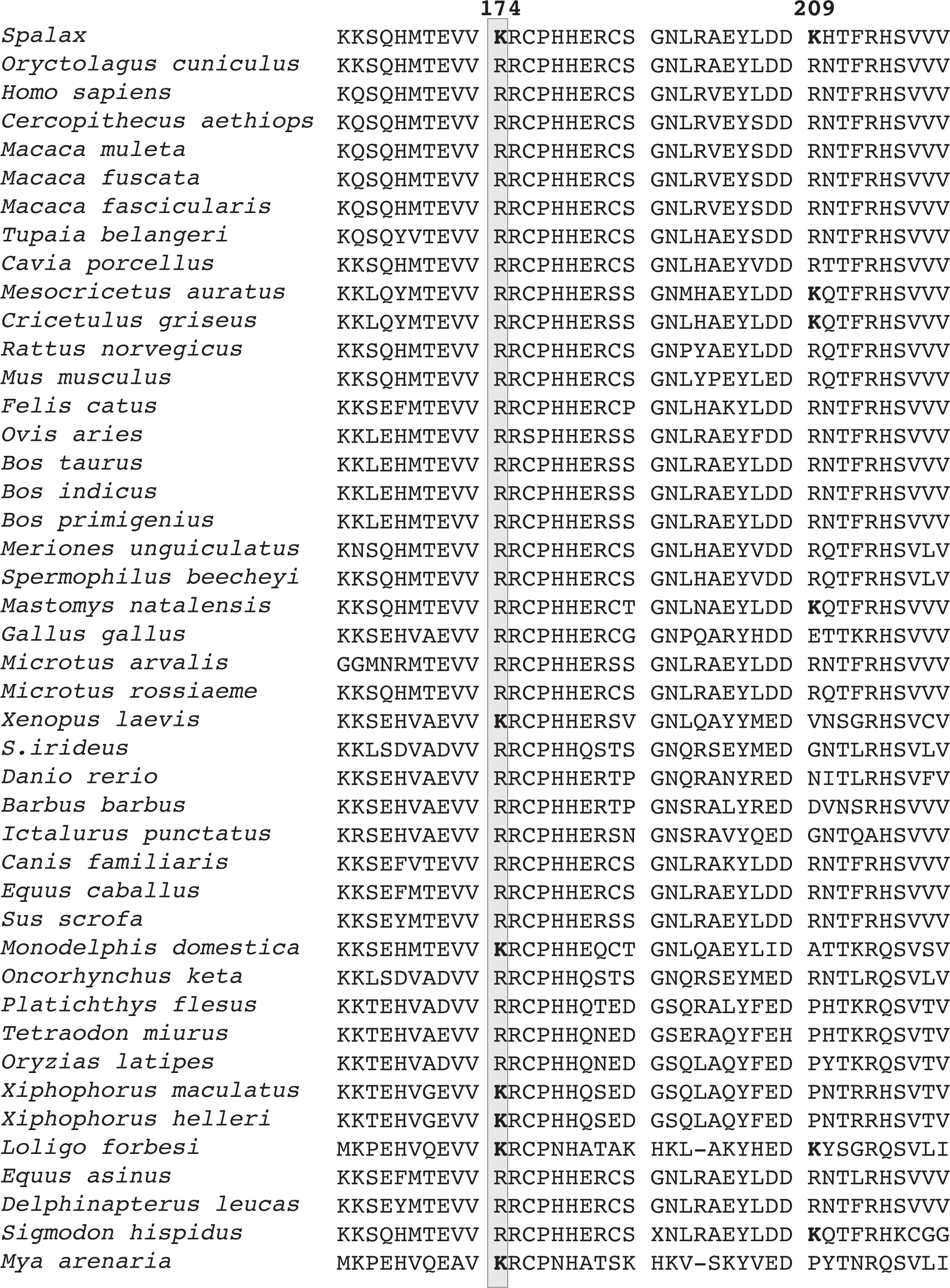
Comparing the p53 DNA binding domain across species. A fragment MSA describing the DNA binding domain, across forty four disparate p53 orthologs, is represented to illustrate the conserved (invariant) amino acid residues and the variants. (Adapted from Reference [117]). **(Inset for Box 2)**

We try to present a broader perspective here, that genome-level molecular adaptations have led to the survival and proliferation of a variety of species in harsh, stressful and disparate environments.

Next, to emphasize further that genomes have evolved uniquely across disparate species, we further synopsize some results from DDR-related pathways (See Box 4).

#### Box 4

##### Adaptations in DDR pathways

Here, using comparative genomic approaches, we highlight recent reports involving various DDR pathways, and encompassing multiple genes. We highlight recent results from the non-homologous end joining (NHEJ) repair pathway and then draw parallels invoking the homologous recombination (HR) repair pathway in bacteria and mammals. Sequence-based computational biology analysis was used to provide insights and perspectives to predict and rationalize adaptive evolution amongst various interacting proteins. First, we synopsize two independent studies from the NHEJ pathway in this context of an adaptive molecular sequence evolution.

An evolutionary screen of proteins from the yeast genome (*Saccharomyces cerevisiae* and *Saccharomyces paradoxus*) identified various NHEJ proteins to be under adaptive evolution [82]. In that study, the authors identified a total of seventy-two positively selected genes from the yeast genome and subsequently focused on pathway-centered analysis where they identified statistically significant positive selection from NHEJ repair pathway genes and genes such as *XRS2, POL4, SAE2* and *NEJ1* [82]. Moreover, they also demonstrated that the identified molecular adaptations are not a result of *Saccharomyces cerevisiae* adapting to laboratory conditions as they obtained similar results by using YJM789, the “wild” *Saccharomyces cerevisiae* isolate [82]. The rapid evolution of those proteins seemed counterintuitive because NHEJ response to DDR is a critical repair response among eukaryotes. In support of their counterintuitive results, the authors hypothesized that those NHEJ genes were under positive selection to counter the integration of Ty LTR-retrotransposons in their genome [82]. In a nutshell, that study primarily identified NHEJ genes manifesting molecular adaptations, in yeast, which were essential for its survival [82].

In an independent comparative genomic study of proteins from the human NHEJ pathway, the authors have delineated human DDR genes under positive selective pressure [120]. Though the critical functions of DNA repair and checkpoint signaling were largely conserved, they identified signatures of positive selection in five NHEJ genes (*XRCC4, NBS1, Artemis, POLλ and CtlP*), during the evolution from simian primates to humans. Interestingly, only one out of nearly seven hundred and fifty codons (<0.5%) of the *NBS1* gene was under positive selection, yet on the other hand they reported *POLλ* having ~5% codons under positive selection, implying that a complicated systems-level genetic underpinnings has been the basis for such diverse quantum of molecular adaptations. In that study, the authors hypothesized that positive selection at select codons might be responsible in providing an evolutionary advantage to the human genome (host genome) from parasitic viral genomes, which was critical for human health and survival [120].

Next, a separate study of human genes from Fanconi anemia *BRCA* (FA/BRCA) pathway is synopsized [121]. That theoretical study investigated a select set of genes: *ATM, BRCA1, BRCA2, CHK2, NBS1, RAD51, FANCA, FANCB, FANCC, FANCD2, FANCE, FANCF, FANCG, FANCL* and *FANCM*, focusing on a pathway-centric approach to identify positive selection in vertebrates [121], about which we suggest an alternate hierarchical analysis that first considered clades of closely related eukaryotes to ensure that on an average *d_N_~l* at the codon level. Using distantly related orthologs, from humans to fish, the author concluded positive selection at specific codons in most of those fifteen genes but negative selection pressure for *RAD51, FANCA* and *FANCG* genes [121].

The divergence of humans from early vertebrates, such as fish, may be traced back to hundreds of million years ago. Ideally, models of codon evolution have been optimized for closely related orthologs, where on the average a single nonsynonymous substitution may be hypothetically anticipated per codon. If judiciously implemented, theoretical approaches from such comparative sequence analysis help identify putative molecular adaptations in a species-specific context, which may subsequently be validated using experimental assays. Using this extensive background, we shall now proceed to appreciate how the genome-level sequence evolution, in the context of human evolution, may help trace the genetic etiology of lifestyle-related complex multifactorial diseases, such as CVD.

## Comparative genomic study can determine a unique hierarchy of common genetic variants from patients with stratified levels of disease severity

Modern genotyping platforms, such as microarrays, enable us to systematically probe the inherited genotype at unique and predetermined SNP loci (for example, the Cardio-MetaboChip from Illumina [122]). An angiography-based study of every CVD patient enabled us to designate a SYNTAX-based score for them and helped us stratify their levels of CVD severity [56, 57] (Fig. 5A). For this comparative genomic analysis, we studied the following cohorts of human subjects: (i) all cases and controls (S_TOT_), (ii) exclusively controls (S_CTRL_), (iii) exclusively CVD patients (S_CVD_), (iv) most severe CVD patients (S_sevCVD1_), (v) CVD patients with intermediate severity (S_sevCVD_2_), and (vi) CVD patients with least severity (S_sevCVD_3_) (Fig. 5A), and probed the genotype of all individuals at SNPs genome-wide. Next, we performed an iterative computational analysis (Fig. 5B), using the genotype from the designated cohorts (Fig. 5A). We first computed the Shannon entropy [123] (See Box 5) at every SNP locus that was probed by our microarray. Furthermore, for quality control purposes, we omitted genotype information recorded by faulty probe(s). After ensuring for that locus, we also computed the zygosity at that locus (Fig. 5B). Independent instances of autosomal recessive [124] as well as dominant mutations [125, 126] have been traced to familial hypercholesterolemia, which in turn associates with CVD. Hence, we sought to compute both homozygous and heterozygous loci that were exclusively invariant among controls and CVD patients.

**Figure 5.**
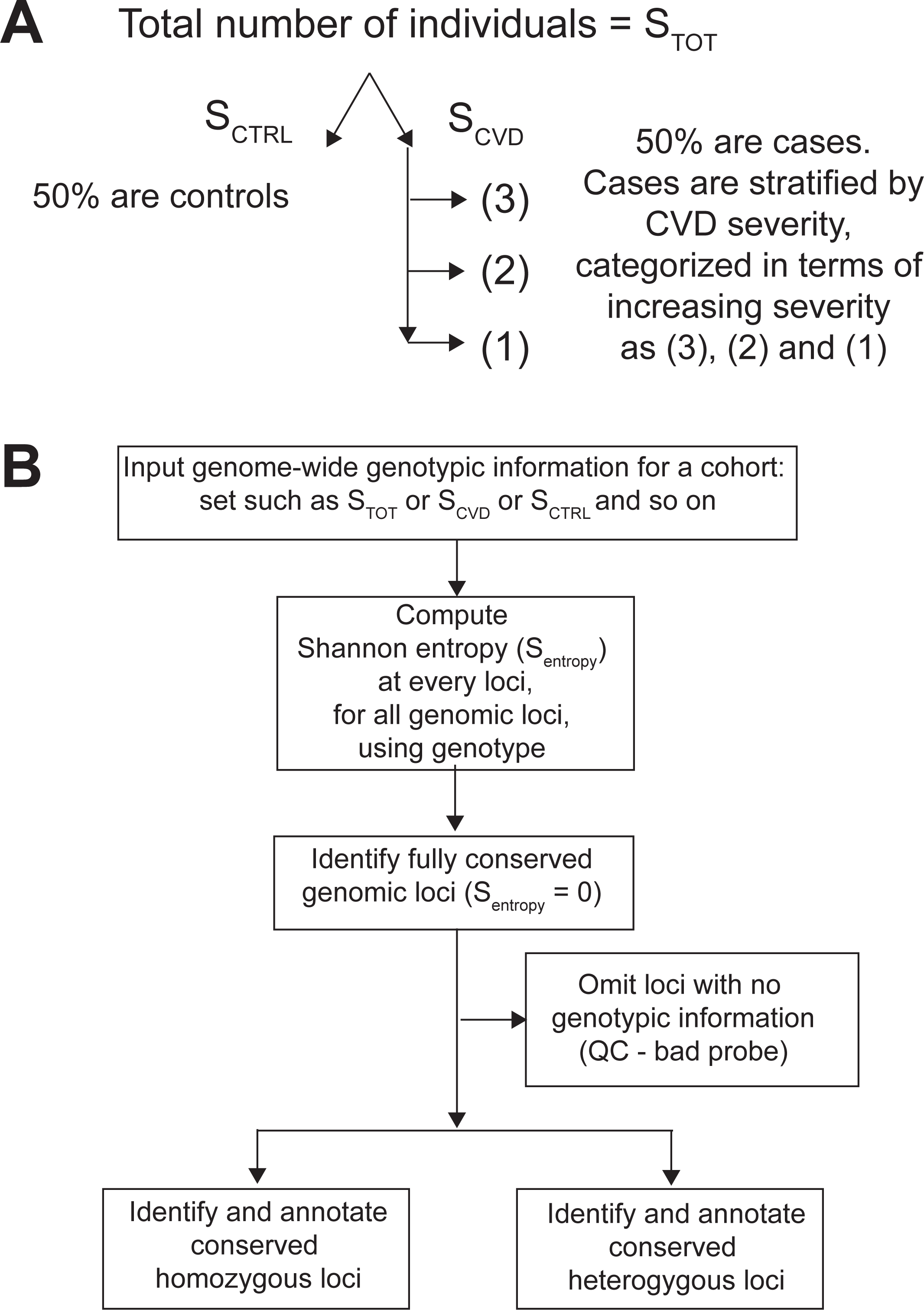
The outline of our bioinformatics analysis used to identify the hierarchy of genetic variants in subjects with disparate levels of disease severity. **A.** The hierarchical organization of data from human subjects used to demonstrate our working hypothesis. **B.** The flowchart describing the iterative comparative genomic analysis. The workflow to describe the computation and quality control steps for identification of invariant genotype across the human genome, for over one million SNP loci. At different stages of analysis, the input dataset for this procedure is different, for example genotypic dataset from (i) case and controls (S_TOT_), (ii) only cases (S_CVD_) (iii) only controls (S_CTRL_), (iv) only severe CVD cases (S_sevCAD1_), (v) only CVD cases with intermediate severity (S_sevCAD_2_), (vi) only CVD cases with low severity (S_sevCAD_3_).

### Box 5

#### Computing the Shannon entropy at the SNP locus

Modern microarray platforms, such as the Cardio-MetaboChip [122], probe the genotypic at predetermined genomic coordinates, which are identical across all subjects – cases and controls. Hence, the genotype data analyzed was analogous to a MSA and ready for a comparative genomic analysis. However, neighboring nucleotides were megabase apart and not adjacent nucleotides from a complete genome. The different genotype probed at a locus, per subject may be annotated in 11 ways. We considered four homozygous calls (AA, TT, GG, CC), six heterozygous calls (AT, AG, AC, TG, TC, GC) and “–” as “bad call” for incorrect / poor quality genomic information that was probed therefore, *N =* 11. Then, for a given study cohort, the Shannon entropy (*S*) at every SNP locus is defined as [123]:

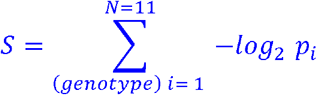

Here, *p_i_* is the probability to find the genotype (out of a total of *N* = 11) at a given locus from an MSA column. The conserved or invariant locus has *p_i_* = 1 for a specific genotype but zero for all other genotypes, hence log_2_ *p_i_* = 0, and therefore Shannon entropy *S* = 0. Once invariant loci are computed, their genotype information is used to identify zygosity at that locus (Fig. 5B).

### CVD cases and corresponding gender- and age-matched controls have a shared ancestry

Although it is common knowledge that the CVD cases and controls from an outbred native subpopulation have a shared ancestry, nevertheless we illustrate this by counting the number of unique invariant genotype for all SNP loci. If individuals have diverged around the same time, from a common ancestor, then the numbers of evolutionarily conserved SNP loci per chromosome may be at par among CVD cases and controls. Such an exercise is motivated because an early onset of CVD is reversible [16] and the human subjects that we studied were all native to India. For a gender- and age-matched case versus control study, the genome-wide identification of statistically consistent number of distinctly invariant loci, on identical chromosomes, may suggest similar evolutionary divergence time from their shared ancestry [36]. We did not do this analysis at the gene-level because most invariant loci were intergenic (unpublished results). Hence, we sought to identify and annotate all invariant loci from the mutually exclusive studies of S_CTRL_ and S_CVD_ respectively. We computed *S* for all SNP loci genome-wide, from a 3-way comparative study, using those two mutually exclusive genomic data sets: S_CTRL_ – S_TOT_ and S_CVD_ – S_TOT_ (Fig. 6A). Here, the subtracted part S_TOT_ implies that all SNP loci that were computed with invariant genotype from the S_TOT_ study were omitted from those in S_CTRL_ and S_CVD_.

**Figure 6.**
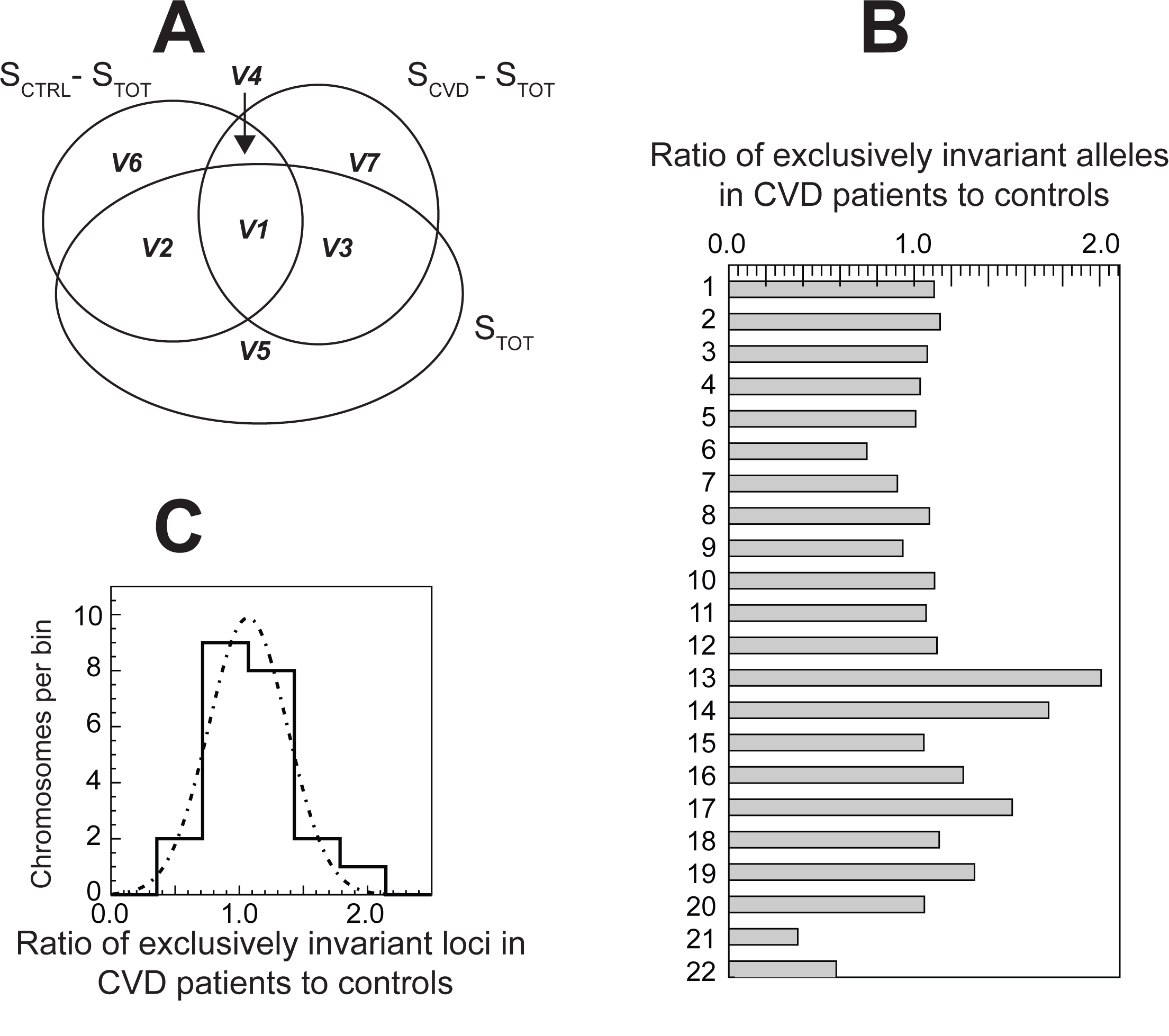
Comparative genomic study demonstrates similar evolutionary divergence times of CVD cases versus age- and gender-matched controls. **(A)** A Venn diagram that represents the study plan for a 3-way comparative genomics approach. **(B)** A bar graph describes the ratio of total number of exclusively invariant loci, apportioned per chromosome, obtained from the CAD cohort with respect to the controls, is illustrated here. All loci that were identical to both cohorts were subtracted first. **(C)** The same ratio from (B) is now described using a histogram, which has been fitted to a Gaussian function (normal distribution).

Here, it is our goal to show that the numbers of exclusively invariant homozygous loci (apportioned per chromosome), and exclusive to either the CVD or control cohorts are at par. Our proposed null hypothesis is that the distribution of all invariant loci from the two mutually exclusive cohorts is statistically consistent. The alternate hypothesis proposed is that the two distributions are statistically inconsistent, suggesting that the two cohorts did not diverge around the same time, from an ancestral population. Using the nonparametric Kruskal-Wallis test to compute if the likelihood of the two distributions was statistically consistent, we report that the null hypothesis is tenable (P < 0.23). In addition, we also computed the ratio of the total number of invariant loci, apportioned per chromosome, from CVD patients with respect to control subjects. We report that the ratio of numbers of invariant loci from the CVD to control cohort is at par (Fig. 6B). On representing the same ratio in a histogram, we report that the histogram mean (Fig. 6C), which confirms that the number of invariant loci from the cases and controls are at par. On repeating the analysis with invariant loci exclusively from the protein-coding genome, we were able to cross check those results and independently computed another histogram with mean, but with an expectedly greater variance because there were fewer invariant loci (results not shown). Taken together, our results support the fact that the individual CVD cases and controls, have diverged from a common ancestral subpopulation and therefore we may attempt to trace the common evolutionary etiology of CVD by tagging loci that were invariant in a hierarchical manner – for example distinct loci were manifest invariant in the combined cohort, among the controls, and among CVD cases.

### 3-way comparative genomic study of invariant homozygous loci

Next, using the previously computed loci, we continued to seek additional insights and perspectives from the study of invariant homozygous loci, which were computed in three different ways: from all individuals, only CVD cases and only controls. To identify and annotate all genes with at least one invariant homozygous locus, which was distinct from the CVD versus control cohort, we did a 3-way comparative genomic analysis of S_TOT_, S_CTRL_ and S_CVD_ cohorts. Using the Venn diagram in Fig. 6A, we represented seven distinct sectors *V1–V7*, each representing the genes or intergenic regions that had such invariant loci from S_TOT_, S_CTRL_, and S_CVD_. We denote those studies as S_V1_–S_V7_ and will only focus on the S_V1_ study to identify genes with the three different types of variants that manifest uniquely in these cohorts. More specifically, we sought genes that represented the sector *V1* from the 3-way overlap (Fig. 6A), which may be represented by hypothetical genes *GENE_G1* and *GENE_G3* (Fig. 7). From this analysis we were able to identify *LDLR* along with other genes (unpublished results). As mentioned earlier, distinct *LDLR* variants are implicated in lipid regulation and metabolism [41], which manifested gain-of-function and loss-of-function, and hence our formalism warrants further scrutiny using this evolutionary-based perspective.

**Figure 7.**
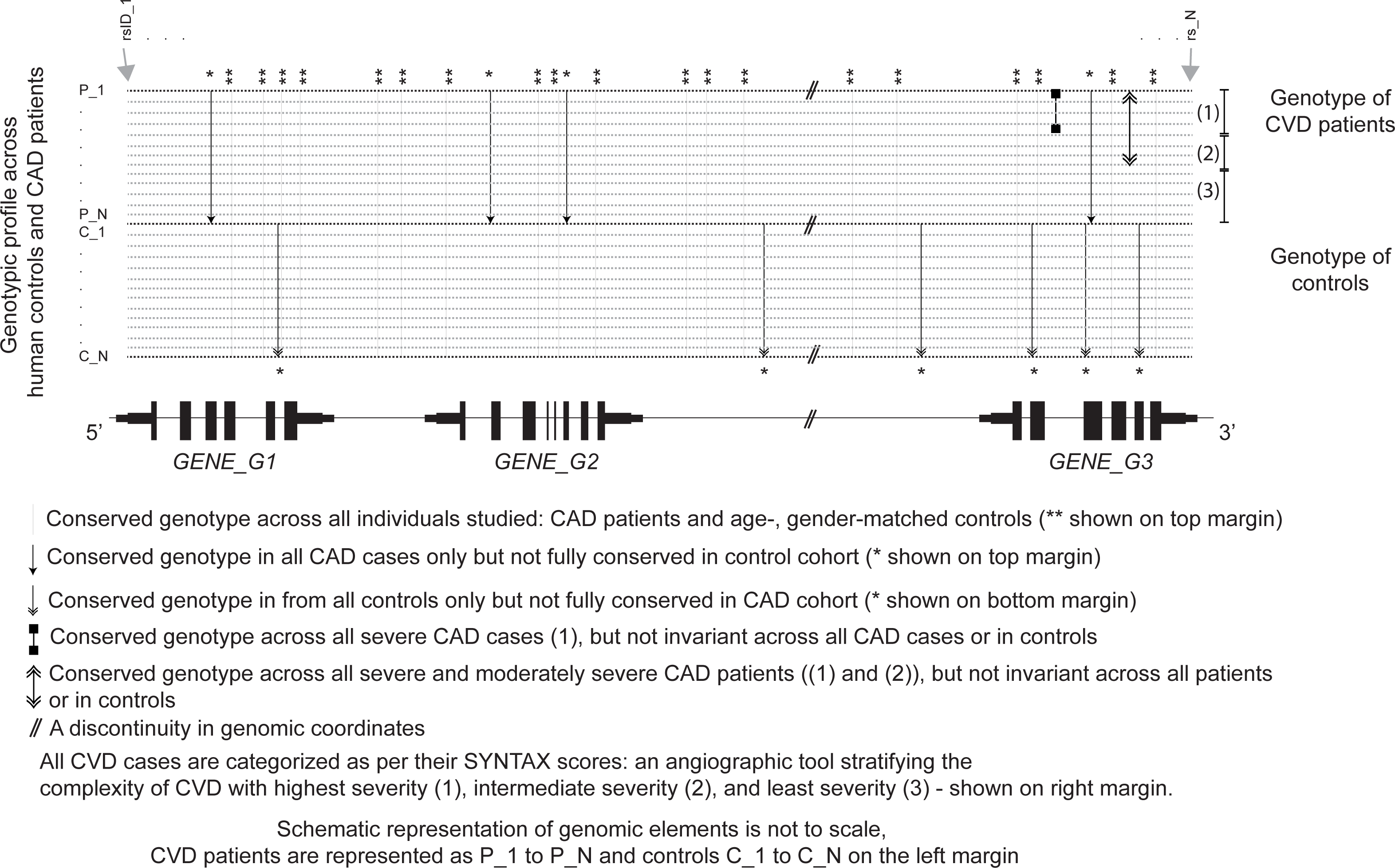
Demonstrating a rational for a hierarchical comparative genomic analysis using hypothetical DNA sequences obtained from cases and controls. Hypothetical DNA sequences obtained from cases and controls for a specific genomic region of interest are illustrated as points. The dotted lines indicate the presence of a string of bases representing the genotype (of a person) at various SNP loci (for example, genome-wide loci probed by a microarray chip). The black rectangles below the sequences reflect hypothetical functional elements (UTRs, exons interspersed by introns) at the positions indicated in the sequence above. A vertical line (grey lines, black arrows with single and double arrowhead) represents the invariant bases (completely conserved genotype) at the given genomic locus. The grey vertical line represents a locus that has invariant genotype in all cases and controls, and with a single arrowhead (double arrowhead) represents exclusively conserved genotype from instances of cardiovascular disease CVD cases (age- and gender-matched controls). A black line with two dots represents systematic and complete genotype conservation at that locus, which exclusively only to severe CVD cases, but not in the remaining cases or controls. The CVD severity among cases is stratified in three tires: (1) highest severity, (2) intermediate severity, and (3) lowest severity, as shown on the right margin.

### Exclusively conserved loci in patients with most severe CVD (S_sevCVD_1_ study)

To further the genetic etiology of CVD, we sought out the common variants that were not conserved in CVD cohort, and with a prototypical hierarchy that represented the paradigm exclusively from the hypothetical *GENE_G3* in Fig. 7. As mentioned before, angiography-based studies (SYNTAX-based scoring [56, 57]) were used to stratify the severity in CVD patients. The patients grouped as most severe (S_sevCVD1_) had multiple manifestations of severe arterial blockages (unreported results). Therefore, to characterize the genetic etiology of severe CVD cases, using the workflow described earlier (Fig. 5B), we sought to compute those invariant homozygous and heterozygous SNP loci that were exclusive only to that cohort (S_sevCVD_1_).

This analysis is analogous to seeking invariant genotype in a MSA, where a few aligned sequences from a set of homologs will yield greater conserved positions, and as more and more homologs are added, the number of conserved positions will recede, or at best remain constant. Hence, we anticipate that the study with most severe CVD patients will result is a larger set of invariant loci than the combined study of patients with least severity, intermediate severity and highest severity. As the cohort size for severe CVD is smaller compared to the full CVD cohort, a genome-wide search for loci with exclusively invariant genotype among them is likely to yield more false positive loci than true positive ones. Therefore, we take a three-tire approach (described next) to sequentially filter out as many false positives as possible, and only compute the homozygous and heterozygous invariant loci that were exclusive to S_sevCVD1_.

### An iterative and hierarchical strategy to identify unique common variants from the cohort of most severe CVD cases (S_sevCVD_1_)

Using S_sevCVD_1_, we compute the for all SNP loci, and delineated the homozygous and heterozygous loci with. In the first tire of analysis, we omitted all invariant loci that were already identified in the S_TOT_ study cohort. Next, in the second tire, using all remaining loci, we subsequently omitted the homozygous / heterozygous invariant loci from the CVD patient study cohort S_CVD_. Finally, in the final third tire, we combined cohorts of intermediate (S_sevCVD_2_) and lowest CVD severity (S_sevCVD_3_) to progressively narrow down our search for invariant SNP loci that were exclusively invariant among the most severe CVD subjects and not otherwise.

In a nutshell, on iterating the analysis outlined here, for different subject cohorts: both case and controls (S_TOT_), only controls (S_CTRL_), only CVD patients (S_CVD_), most severe CVD patients (S_sevCVD_1_), CVD patients with intermediate severity (S_sevCVD_2_) and CVD patients with least severity (S_sevCVD_3_), and filtering out SNV loci obtained from any cohort at higher hierarchy, we will systematically gain access to the genetically invariant loci for cohorts such as S_sevCVD1_, S_sevCVD_2_ and S_sevCVD_3_.Therefore, using a 3-way and three-tire severity-derived CVD data, a robust comparative genomics study is proposed to understand why common genetic variants persist as invariant loci only in very specific cohorts, such as the most severe CVD cohort, and why the same genes manifest markedly different and hierarchical variants.

Moreover, after classifying CVD patients by their levels of severity, a comparative genomic analysis across the severe category will surely enable us to identify the invariant loci that are unique among identical set of genes or intergenic regions (*GENE_G3* in Fig. 7). We report that such invariant loci are largely inter-genic, with consecutive invariant loci span genomic distances in megabase scale. However, it is beyond our scope to determine if those variants represent a gain-of-function / loss-of-function paradigm [48, 49], or are benign [127].

Genes computed using this 3-way comparative genomic study represent putative candidates for multifactorial CVD severity. Therefore, we hypothesize here that genes that manifest evolutionary plasticity from: (i) the 3-way analysis study of S_V1_, (ii) with exclusively invariant homozygous and/or (iii) heterozygous loci in the severe CVD cohorts, may be used to trace the hierarchical etiology of this complex multifactorial disease, either directly modulating physiological function or via its interactors, so as to reverse the disease prognosis [16]. We also emphasize that our results are along the lines of the seminal discovery involving evolutionary plasticity of the *LDLR* gene, wherein it was shown that unique variants of the gene gave rise to both loss-of-function and gain-of-function [41]. Hence, more importantly a candidate-gene based search strategy may putatively enable us to identify common genetic variants having unique evolutionary plasticity that is manifest in diseases with differential severity.

## Conclusion

In this perspective, we have presented alternate perspectives and insights to trace the etiology of CVD from a very small study set (50% CVD cases and 50% controls) representing an outbred subpopulation from India. We show that it is possible to identify genetic variants from CVD patients stratified by disease severity. The hierarchical plasticity of those variants led us to identify a constellation of genetic variants unique to CVD cases that were stratified by differential levels of disease severity. Consequently, interactions among various proteins that originate from variant genes may give rise to a differential response. Taken together, this work opens up avenues for inter-disciplinary analyses that may complement genome wide association studies. As a primer to computational molecular evolution, we have synopsized that genomes have evolved uniquely in among disparate species, from independently published reports. Accordingly, we highlighted instances of diverse molecular adaptations among interacting proteins across species. Novel insights and perspectives developed here are for motivating a paradigm shift and for wider education that may lead to a better understanding of disease paradigm using an evolution-based approach.

## Outlook

The perspective presented is not restricted to CVD or lifestyle-related diseases and may be explored in the context of diverse diseases, for example autism where in a rational basis for stratification of disease severity exists.

**Table 1.**
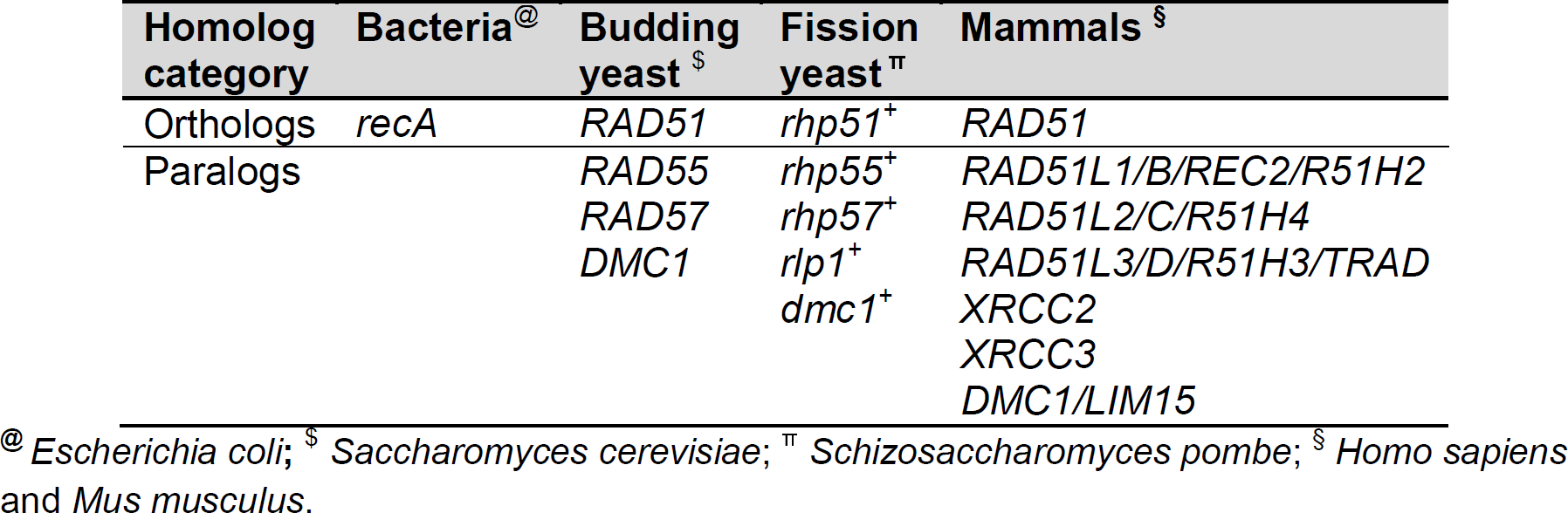
The paralogs of *RAD51* human DNA repair genes and their orthologs from bacteria (*recA* genes) and yeast are illustrated. (Adapted from Reference [86]).

## Acknowledgements

SNF would like to acknowledge Tata Institute of Fundamental Research (TIFR)-DAE, Government of India and Professor BJ Rao of TIFR, Mumbai for financial support. Doctor TF Ashavaid, P.D. Hinduja Hospital and Medical Resarch Centre, Mumbai for sharing a microarray-based genomic data from 149 CVD cases and 149 controls.

